# Non-invasive vagus nerve stimulation in a hungry state decreases heart rate variability and wanting of a palatable drink

**DOI:** 10.1101/2022.05.18.492424

**Authors:** Zeynep Altınkaya, Lina Öztürk, İlkim Büyükgüdük, Hüseyin Yanık, Dilan Deniz Yılmaz, Berçem Yar, Evren Değirmenci, Uğur Dal, Maria Geraldine Veldhuizen

**Author notes:** Corresponding author: Maria Geraldine Veldhuizen.

## Abstract

Vagus nerve signals from the gut to brain carry information about nutrients and drive food reward. Such signals are disrupted by consuming large amounts of high-calorie foods, necessitating greater food intake to elicit a similar neural response. Non-invasive vagus nerve stimulation (nVNS) via a branch innervating the ear is a candidate treatment for obesity in humans. There is disagreement on the optimal location of nVNS in the ear for experimental and clinical studies. There are also no studies comparing nVNS in hungry and full states. We aimed to compare ear position(s) for nVNS and explore the effects of nVNS during hungry and full states on proxies for autonomic outflow (heart-rate variability) and efferent metabolism (gastric frequency and resting energy expenditure).

In a within-subject design, 14 participants (10 women, on average 29.4 +/- 6.7 years old) received nVNS in four different locations (cymba conchae, tragus, earlobe, or tragus AND cymba conchae) on separate days. In each session, participants were asked to consume a palatable chocolate flavored milk. With electrography on the abdomen and indirect calorimetry in a canopy, we measured electro-cardiogram, electro-gastrogram and resting energy expenditure for 15 minutes before and at least 35 minutes after consumption of the palatable drink. We also collected ratings of the palatable drink and internal and other states.

Pre-drink consumption (in a hungry state) we observed no differences in the effect of location of acute nVNS on resting energy expenditure and gastric frequency. However, nVNS in cymba conchae decreases heart-rate variability and ratings of how much participants want to consume the drink. After drink consumption and with continued nVNS, gastric frequency is unchanged, and resting energy expenditure increases regardless of stimulation location. Heart-rate variability decreases in all locations, except cymba conchae. We also observe a trend for an increase in gastric frequency in late post-drink consumption time-points in cymba conchae.

These results suggest that nVNS in the cymba conchae in a hungry state has a similar acute effect on vagal tone as food consumption: to decrease heart rate variability. This effect then negates the usual postprandial effects of a decrease in heart rate variability as seen in the other nVNS locations. This suggests that nVNS in cymba conchae may act primarily on vagal afferent autonomic (and only modestly on metabolic output) in a similar way as food consumption does.

**Highlights:**

- We measured autonomic outflow and efferent metabolism before and after consumption
- We manipulated the location of nVNS stimulation in the outer ear
- The different locations were earlobe, cymba conchae, tragus, cymba conchae+tragus
- nVNS in cymba conchae decreases <u>pre</u>-consumption heart-rate variability and wanting
- nVNS in other locations decreases <u>post</u>-consumption heart-rate variabilty

## Introduction

The vagus nerve (VN) carries signals about food from gut to brain [1–3]. Both information about volume (from mechano-receptors [1]) and nutrients (from sensory neurons sensitive to metabolic signals [4, 5]) are transmitted through this route. Opto- and chemogenetic experiments showed that vagal sensory neurons drive post-ingestive fat [6] and sweet mediated reward [7]. These pre-clinical findings reveal a gut-to-brain post-ingestive fat and sugar-sensing pathway critical for the development of food preference. If these data translate to humans, these findings would support targeting the VN to modulate food preferences.

Vagal nerve stimulation (VNS) involves implanting electrodes on the cervical vagus nerve [8]. Some retrospective studies in humans treated for epilepsy or depression show significant weight loss in participants [9, 10], and in a prospective study a decrease in the preference for sweet food images was observed [11]. Non-invasive VNS via the auricular branch (transcutaneous non-invasive VNS, nVNS) [12, 13] is effective in treating depression in clinical studies [14], and has enabled experimental studies in healthy human participants [15, 16]. The optimal location in the outer ear for nVNS is contested, as both tragus and cymba conchae have vagal innervation [17,18]. The preferred choice for experimental studies is the cymba conchae due to the exclusive innervation by the vagus, as influences from other auricular nerves can be ruled out. However, from a clinical perspective this may not be the optimal choice, as effectiveness may be greatest from stimulation of all vagally innervated auricular regions. To our awareness, no studies have examined the effectiveness of nVNS in a combination of vagally innervated auricular areas.

Recent works using nVNS modulation of neuro-behavioral and physiological outcomes relevant to food reward and metabolism in humans show mixed results. Various recent studies examining neuro-behavioral outcomes have used food stimuli or tasks related to food reward. Application of nVNS in cymba conchae immediately prior to a test session had no effect on electro-encephalogram responses to visual food stimuli relative to other objects, as well as no effect on food intake [19]. Neural responses measured with fMRI were stronger to liked food images in food reward regions (midbrain, thalamus, hippocampus, amygdala and medial orbitofrontal cortex) relative to food images that received lower hedonic ratings after nVNS in cymba conchae [20]. Liking and wanting ratings of food images were not affected in healthy controls, but in participants with major depressive disorder liking ratings increased under concurrent nVNS in cymba conchae. We previously showed an effect of concurrent nVNS in cymba conchae on liking of orally sampled foods, such that liking of low-fat puddings, but not of high-fat puddings, was increased [21]. Food intake was not affected in this study. nVNS in cymba conchae also increased participants’ drive to obtain less-wanted prospective food rewards in an effort allocation task [22]. Lastly, two weeks of nVNS in tragus concurrent with bottle-feeding improved oral intake in about half of premature or brain injured infants who had failed oral feeding until that time [23].

Similarly, inconsistent effects were observed for the effect of nVNS on physiological outcomes related to metabolism of food. In the absence of food consumption, nVNS in cymba conchae decreases gastric frequency and has no effect on resting energy expenditure [24]. Post consumption of a glucose drink nVNS in cymba conchae did not affect resting energy expenditure, or blood metabolites [25].

The studies that focused on neuro-behavioral and physiological outcomes relevant to food reward and metabolism all examined the effect of nVNS by comparing sham stimulation in the ear lobe to verum stimulation in the tragus <u>or</u> cymba conchae, but to our awareness none examined effects of combined stimulation in the cymba conchae <u>and</u> tragus. The studies that examined the effects of nVNS on proxies of efferent vagal effects relevant to food metabolism generally did so in the absence of food consumption [24] or with a model stimulus (glucose drink,[25]), without explicitly looking at effects in a hungry vs a full state.

Here we use a protocol with a naturalistic food stimulus; a palatable chocolate-flavored milk. We evaluate the effectiveness of nVNS with three physiological proxies for vagal tone: autonomic outflow through heart-rate variability, and efferent metabolism through electrogastrogram and resting energy expenditure. We measured the physiological outcomes variables in a 15-minute pre-drink (hungry state) period and then in a 35-minute post-drink (full state) period. nVNS stimulation was continuous throughout this period, but on separate test days we stimulated different locations of the ear: ear lobe (sham/placebo control condition), and verum stimulation in either cymba conchae, tragus, or in both cymba conchae *and* tragus.

We first predicted that if nVNS modulates food reward processing in humans, we should observe differences between the stimulation locations that have vagus nerve innervation (cymba conchae, tragus and cymba conchae + tragus) versus the sham location (ear lobe) on proxies for autonomic outflow and efferent metabolism after consumption of a palatable drink (in a full state), but not before consumption of a palatable drink (in a hungry state).

Our second prediction was that if the vagally innervated areas of the outer ear in cymba conchae and tragus are functionally equivalent, we should observe that stimulation of the combination of cymba conchae + tragus leads to effects on the proxies for autonomic outflow and efferent metabolism that are in the same *direction* (relative to sham in ear lobe) as the separate locations in cymba conchae or tragus. We also predicted that the effect of the combination of stimulation in cymba conchae + tragus would have a *magnitude* that is equal or greater than the separate locations in cymba conchae or tragus, but not smaller.

## Material and methods

### Participants

Fifteen healthy, participants [11 women, 4 men] with a mean (± standard deviation) age of 28.9 (± 6.8) years [range: 21-43], with a mean body mass index (BMI) of 23.5 ± 3.2 kg/m^2^ [range: 18.6-29.3] and body fat percentage (measured with Tanita BC 601, Tanita Corporation, Tokio, Japan) of 26.9 ± 5.2 [range: 18.6-42.8] participated in the study. All participants reported having no known taste, smell, neurological, psychiatric (including eating disorders), cardiological, metabolic or other pathological disorders. The data for this study were collected between May and August of 2021. During this period Mersin University was encouraging working from home and all teaching was remote. Therefore four participants were part of the research team. Other participants were recruited by word of mouth from around campus by the research team. The Mersin University Committee for Clinical Research approved the study protocol (number 7807789 / 050.01.04 / 1152384) and written informed consent was obtained from all study participants. All participants reported having not had COVID-19 and most participants had been vaccinated with two doses of Sinovac. The data from one participant was excluded due to technical difficulties with the nVNS stimulation during one of the sessions. The remaining 14 participants [10 women, 4 men] were 29.4 (± 6.7) years old, had a BMI of 23.8 ± 3.2 kg/m^2^ and body fat percentage of 27.0 ± 5.4.

### Design & session procedure

We used a within-participants design in which all participants were exposed to the sham condition and other nVNS stimulation locations on separate days, in a counterbalanced order. Participants were scheduled for the same time of day for each session, usually in the morning. Sessions were scheduled in the follicular phase or luteal phase for participants with a menstrual cycle. We used a within-participants design, with 4 sessions per subject. During each session nVNS was delivered to a different location: cymba conchae, tragus, lobe, or tragus <u>and</u> cymba conchae (counterbalanced across participants). The session was completed within approximately 90 minutes for each participant. The four sessions of each participant were usually completed within a week, and a maximum of 2 weeks. Instructions we gave participants before session; avoid heavy exercise the day before, should not consume alcohol the day before, coming to the test fasted for at least 4-5 hours, consume similar meals before their fast on each test day, no caffeine consumption up to 4 hours before and should not consume nicotine/cigarettes for at least 2 hours before the test. These instructions comply with best practices for the measurement of both resting energy expenditure and heart-rate variability [26–28]. The day before each session they were contacted to inquire about COVID-19 symptoms and sessions were rescheduled if any symptoms were reported.

Each session included the same events in the same order (Figure 1C). All sessions were performed in a thermoneutral, quiet and semi-dimmed room. The room temperature was 20-24°C and humidity was around 50%. First, participants were asked to void their bladder and rest for 15 minutes. Then they were outfitted with the electrodes for electro-cardiogram (EKG) and electro-gastrogram (EGG) measurement and tVNS stimulation. Then we adjusted tVNS stimulation intensity for each participant individually (see nVNS section below for details). Then we asked participants to rate the perception of the nVNS stimulation on their ear and their internal and other states (see detailed descriptions below). Then the participant laid down on a semi-reclined hospital bed. To measure resting energy expenditure (REE), a canopy hood was placed over the upper body of participants with a veil to prevent any air leakage while participants were lying in a semirecumbent position. Disposable antiviral filters were used for each participant in all sessions as a precaution for COVID-19. We asked participants not to sleep and move as little as possible. We measured the EKG, EGG and REE simultaneously for 15 minutes. Then we gave a palatable drink to participants after the top of the bed was lifted. They took a sip first with a pipette without moving much and then they made the drink perception ratings. They then consumed the rest of the drink and made the internal state and other ratings. We continued to measure energy expenditure after consumption of the palatable drink for at least 35 minutes. Participants rated the perception of the nVNS stimulation on their ear and their internal and other states at the end of the session.

**Figure 1.**
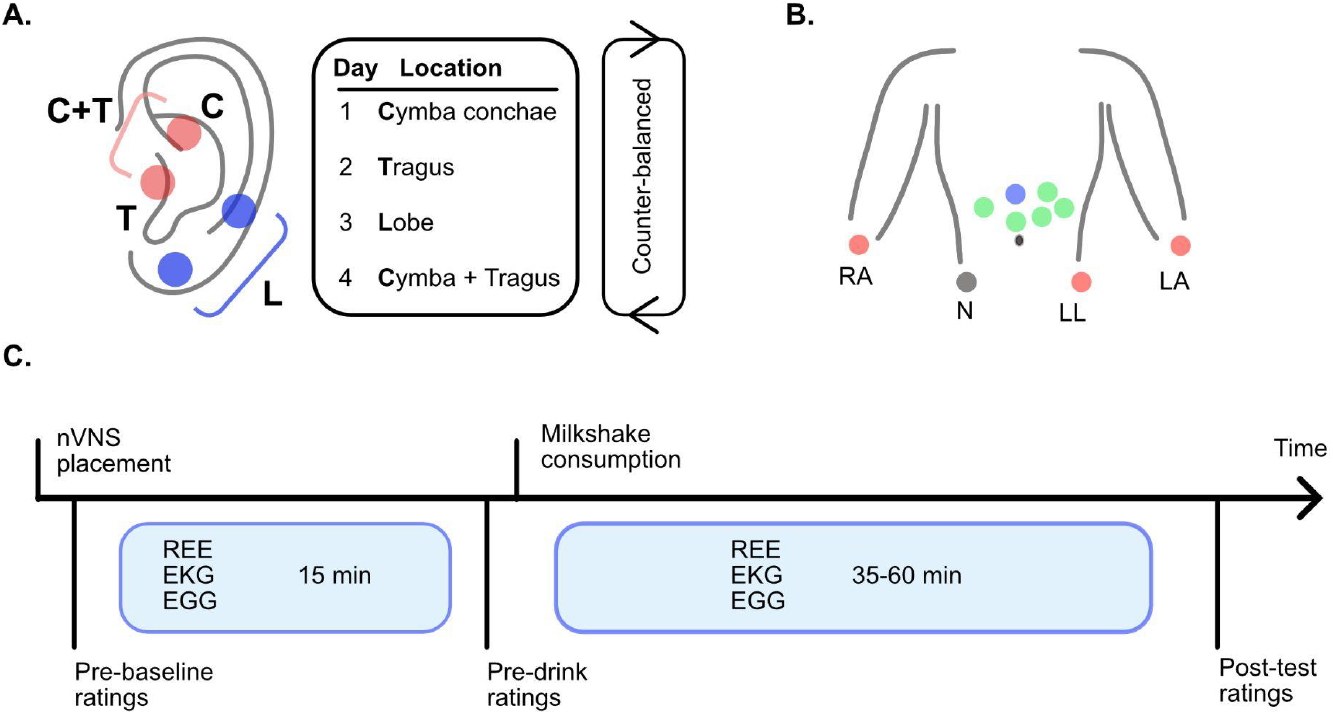
Graphical overview of the electrode locations and test day protocol. A. Ear locations for nVNS (non-invasive vagus nerve stimulation). 4 conditions for ear stimulation as Cymba (C), Tragus (T), Lobe (L), Cymba + Tragus (C+T) applied counterbalanced during four test days. B. Electrode locations for electrogastrography (EGG) and electrocardiography (EKG) recordings during resting energy expenditure (REE) measurements. C. Test day protocol. Before measurements, the earpiece was placed according to the ear location that is going to be stimulated. After the participants finished the pre-baseline ratings, 15 min of REE measurement was recorded simultaneously with EGG and EKG. Before milkshake consumption, pre-drink ratings were collected. After the milkshake was consumed, 35-60 min of REE measurement with EGG and EKG was recorded. Lastly, post-test ratings were collected.

### nVNS stimulation

Mild transcutaneous electrical stimulation was applied to ear lobe (sham/placebo control condition), and verum stimulation in either cymba conchae, tragus, or in both cymba conchae *and* tragus on separate visits (order counterbalanced over participants). We used commercially available TENS (transcutaneous electrical nerve stimulator, Twin Stim^®^ Plus 3rd edition, Roscoe Medical Inc.) or Vagustim (Vagustim Sağlık Teknolojileri A.Ş., Istanbul, Turkey) stimulation devices. For all locations we stimulated both the left and right ear. For the sham stimulation, we used standard nVNS ear clips with electrodes. Stimulation points were selected in the superior scapha (midpoint of the outer ear margin) and earlobe, where there is no vagus nerve distribution. After one of the two unilateral electrodes was placed on the earlobe, one electrode in the opposite direction was placed on the superior scapha [29]. For the cymba conchae we used electrodes that are mounted on a round plastic stabilizer that fits into the cavum concha (similar to the Cerbomed Nemos^®^ device). In the tragus condition, double sided-clips were clamped onto the inner and outer surfaces of the tragus. For the combined stimulation of both cymba conchae *and* tragus two electrodes in opposite directions were placed inside the cavum concha facing the cymba auricularis and the inner surface of the tragus (see Figure 1A). Most participants were blinded to the sham/verum status of the stimulation locations, however, the research team was not, and four of the participants were part of the research team. For all locations, nVNS stimulation used the following parameters: a biphasic square wave pulse at 25 Hz and a pulse width of 250 μs, with a duty cycle of 30 s on, 30 s off. The total stimulation duration was approximately 90 minutes, the time needed to complete the baseline and post-drink recording periods. Stimulation was applied with constant voltage. These parameters are reported in agreement with proposed reporting guidelines [30]. The amplitude of stimulation was calibrated for each session (stimulation location) and participant individually with a procedure commonly used [30,31], intended to adjust the amplitude to the highest stimulation level that can be reached without causing pain or discomfort. Since there are tissue and innervation differences in the sham and nVNS locations, the resulting stimulation amplitudes may differ between sites, but the calibration ensures that the sensation between sites remains comparable, thus controlling for perception-related placebo effects. During the adjustment procedure, the stimulation was gradually increased. The participant was asked to report when a “pricking, stinging or burning” sensation was felt, which indicated their pain threshold. The stimulus intensity was then immediately decreased gradually until the participant reported an innocuous, comfortable “tingling, vibrating or drumming” sensation. The intensity of the stimulus remained at that selected level for the duration of the session unless the participant reported discomfort, in which case, the stimulus intensity was decreased in the same manner as during calibration to relieve discomfort. For 10 out of a total of 56 sessions, we reduced the intensity later in the session. To confirm iso-intense stimulation, participants rated the sensation of nVNS stimulation on the ear (described in detail below). This type of stimulation is safe and well tolerated [32]. No adverse events were observed.

### Internal and other state ratings

Participants were asked to rate their internal and other states on horizontal 101-point Visual Analog Scales (VAS), presented on a monitor with a dark background, by dragging a cursor across a line. They made ratings of the following states in this order: hunger, fullness, thirst, need to pee, stress, comfort, sleepiness, tiredness, temperature, canopy temperature and canopy airflow. Hunger, fullness, thirst, need to pee, stress, comfort, sleepiness, and tiredness were rated on a unipolar 0-100 VAS with two anchors, labeled with “not at all […]” on the left anchor and “extremely […]” on the right anchor. Temperature, canopy temperature and canopy airflow were rated on a −50 - 50 bipolar VAS, labeled with “very cold/airless” on the left anchor, “good” on the middle anchor and label “very hot/windy” on the right anchor. Each VAS (in this section and below was explained upon the first presentation during the first session to the participant. These and other ratings were collected to address potential confounding variables, but also to monitor participants’ (dis)comfort during the session and make adjustments if necessary.

### nVNS stimulation perception ratings

Participants were asked to rate the sensation of the nVNS stimulation on the ear via unipolar VAS scales for the following sensations in the following order: intensity, tingling, vibration,itching, stinging, and burning (left anchor labeled with “no sensation” and right anchor labeled with “most intense sensation”). Participants have also rated the pleasantness of the nVNS sensation via a bipolar VAS with “very unpleasant” on the left anchor, “neutral” on the middle anchor and “very pleasant” on the right anchor.

### Palatable drink

The palatable drink was a chocolate flavored milk, made from commercially available ingredients. The flavored milk was prepared with 10% fat and 10% sugar weight-by-weight (w/w). The ingredients used were 3.0% fat shelf-stable milk (SEK), 35% fat shelf-stable cream (İçim), white sugar (Balküpü), vanilla sugar (Dr. Oetker), cacao powder (Dr. Oetker Gourmet Cacao). We mixed the chocolate milk one day before the test day and kept it in the refrigerator after waiting for it to cool to room temperature. On the test day, the chocolate milk was taken out of the refrigerator, stirred, and brought to a temperature of 37°C, before being served to the participant in a 200 gram portion (~290 kcal) with an opaque container with a flexible straw.

### Palatable drink perception ratings

Participants were instructed to take a single sip of the palatable drink, swallow it and then make ratings of the following attributes in this order: intensity, liking, familiarity, sweetness, sourness, bitterness, creamy, fatty, other, caloric-ness, healthiness, and desire to consume more of the drink (referred as “wanting” from here on). Liking and familiarity were presented as a bipolar VAS with “not liked/familiar at all” on the left anchor, “neutral” on the middle anchor and “very liked/familiar “ on the right anchor. Healthy was also presented as a bipolar VAS with “unhealthy” on the left anchor, “neutral” on the middle anchor and “very healthy” on the right anchor. Intensity, sweetness, sourness, bitterness, creamy, fatty, and other were presented as a unipolar VAS labeled with “not at all […]” on the left anchor and “extremely […]” on the right anchor. Lastly, “caloric-ness” was presented on a unipolar scale with “zero calories” on the left anchor and “very caloric” on the right anchor”.

### Electro-cardiogram/electro-gastrogram

We used the Kardinero EKG Master USB device and WinEKG Pro software (Kardinero Medikal Sistemler, Ankara, Turkey) to record cardiac and gastric electrical activity by means of standard snap-type cutaneous and disposable electrodes placed on the abdomen. The Kardinero is a standard 12-lead ECG system with a maximum sampling rate of 4000 Hz. We placed four electrodes on limb extremities in the augmented unipolar limb leads configuration: left arm (LA), right arm (RA), left leg (LL) and right leg (RL, neutral) for recording Einthoven’s EKG derivations, and the remaining 6 electrodes were placed relative to anatomical landmarks of the umbilicus, xiphoid process and clavicle as described in Wolpert et al. [33] (Figure 1B). EKG and EGG data were measured simultaneously and both have a sampling rate of 1000 Hz. For heart-rate variability data, we used the recordings from Lead II derivation among the 12-lead EKG, since Lead II direction is similar to the route of the electrical activity of heart [34].

### Indirect calorimetry

REE was measured by indirect calorimetry with the Quark RMR device and OMNIA Cardiopulmonary Diagnostic Software (Cosmed, Rome, Italy). Quark RMR is an open-circuit calorimetry system and equipped with a canopy hood that allows the participant to breathe comfortably. The turbine flowmeter of the Quark RMR device measures the flow rate directly. Before each test session, calibration of flowmeter was performed with a 3 L calibration syringe and gas calibration was done by using certified standard gas tubes containing 16% O2, 5% CO2 and Balance N2. Gas pressure was kept above 10 bar. O2 consumption (VO_2_) and CO2 production (VCO2) were recorded every 10 seconds. We will refer to energy expenditure as resting energy expenditure (REE) regardless of time (pre/post consumption).

### Data-preprocessing general

For EKG and EGG, we divided the entire 15-minute pre-drink time interval into three 5-minute bins. 5 minutes is the recommended gold-standard duration for heart-rate variability recordings [26] and at an expected frequency of three gastric cycles per minute, 5 minutes also allow a relatively stable estimate for EGG [33]. We then averaged the calculated measures (see below) across the three pre-drink time bins to obtain a single estimate of the pre-drink measurement. Best practices for baseline REE data analysis require the identification of a time-interval with a coefficient of variation below 5% [28]. For all three physiological measures, we have divided the post-drink time interval into 5-minute bins. The post-drink time interval was variable in duration, as some participants needed to end the session faster due to discomfort than others. We were able to collect a total of seven 5-minute bins for all participants for all physiological outcome measures.

### Data-preprocessing heart-rate variability

We used Matlab to scale the data to the maximum value and extract baseline drift using a high pass filter with a cut-off frequency of 0.5 Hz. We also applied a bandstop filter to eliminate 50 Hz power line interference. We then used the R package rsleep [35] to extract time of each heart beat using the QRS complex based on the method by Pan & Tompkins [36]. Then we used the R package rHRV [37] to calculate the root mean square of successive differences (rMSSD) in each 5-minute bin pre- (three bins) and post-drink consumption (seven bins). rMSSD is the recommended [26] and most commonly [38] evaluated measure of heart-rate variability in nVNS studies.

### Data-preprocessing electro-gastrogram

We used the Matlab toolboxes Fieldtrip [39] and electro-gastrogram scripts [33] to calculate the electro-gastrogram frequency and amplitude. The algorithm filters the signal and out of the electrodes 1-6 on the abdomen, it determines the optimal electrode based on the peak frequency in the expected range for electrogastrogram. Then, we calculated the mean gastric frequency (MGF) in mHz and amplitude (MGA) in μV for three 5-minute pre-consumption bins and each of the seven 5-minute bins post-drink consumption.

### Data-preprocessing resting energy expenditure

We used in-house written R functions to pre-process the raw resting energy expenditure (kcal/day), calculated from the abbreviated Weir equation [40] from VO_2_ and VCO_2_ measured in every 10 seconds. We confirmed a stable 5-minute measurement with a coefficient of variation in VO_2_ of less than 5% [28], which was used as the pre-drink baseline bin. We then binned across 30 measurements in each of the seven 5-minute bin of post-drink consumption to obtain an average estimate of REE.

### Statistical analysis for the nVNS stimulation settings

We conducted a repeated-measures ANOVA with within-subject factor location (sham, cymba conchae “C”, tragus “T” and cymba conchae and tragus “CT”) on the selected intensity setting (amplitude) of the nVNS stimulation. Planned follow-up t-tests were conducted to compare the effect of location (Bonferroni-corrected for 6 comparisons). Alpha was set at 0.05 here and in all following analyses.

### Statistical analysis for the nVNS perceptual ratings

We conducted a repeated-measures ANOVA with within-subject factor location (sham, cymba conchae “C”, tragus “T” and cymba conchae and tragus “CT”) on each of the sensations that was rated on the VAS scales; intensity, pleasantness, burning, stinging, itching, tingling and vibration. Planned follow-up t-tests were conducted to compare the effect of location (Bonferroni-corrected for 6 comparisons).

### Statistical analysis for the internal and other state ratings

For each of the rated internal and other state ratings on VAS scales (hunger, fullness, thirst, need to pee, stress, comfort, sleepiness, tiredness, temperature, canopy temperature and canopy airflow), we conducted a repeated-measures ANOVA with within-subjects factors time (pre-session, post-drink and post session) and location (sham, cymba conchae “C”, tragus “T” and cymba conchae and tragus “CT”). Planned follow-up t-tests were conducted to compare the change from pre-session to post-drink and from pre-session to post-session for all nVNS stimulation locations together and for each nVNS stimulation location separately (Bonferroni-corrected for 2 comparisons). We also conducted planned follow-up t-tests to assess whether the change from pre-session to post-drink differed between nVNS stimulation locations (Bonferroni-corrected for 6 comparisons). Last, we conducted planned follow-up t-tests for the effect of nVNS stimulation location (Bonferroni-corrected for 6 comparisons) at each of the time-points separately.

### Statistical analysis for the physiological outcome measures

For each of the physiological outcome measure we conducted a repeated-measures ANOVA with within-subjects factors bin (pre-drink and post-drink bins 1-7, corresponding to minutes 1-5, 6-10, 11-15, 16-20, 21-25, 26-30 and 31-35 respectively) and location (sham, cymba conchae “C”, tragus “T” and cymba conchae and tragus “CT”). Planned follow-up t-tests were conducted to compare effects of location (Bonferroni-corrected for 6 comparisons) in each of the bins. We also conducted planned follow-up t-tests to assess the effect of consuming the drink for each of the post-drink bins relative to the pre-drink bin (Bonferroni-corrected for 7 comparisons). We also compared the change from pre- to post-drink consumption for each nVNS stimulation location and performed one-sample t-tests to test against 0.

### Statistical analysis for the palatable drink ratings

We conducted a repeated-measures ANOVA with within-subject factor location (sham, cymba conchae “C”, tragus “T” and cymba conchae and tragus “CT”) on each of the hedonic and sensation of the palatable drinks that was rated on the VAS scales; intensity, liking, familiarity, sweetness, sourness, bitterness, creamy, fatty, other, caloric-ness, healthiness, and wanting. Planned follow-up t-tests were conducted to compare the effect of nVNS stimulation location (Bonferroni-corrected for 6 comparisons).

## Results

### nVNS is not different in perceived intensity, pleasurableness across locations

After placement of the electrodes, the intensity (amplitude) of stimulation was adjusted to the perceptual target of “as intense as possible without getting uncomfortable” (see Methods) and the amplitude setting recorded. The adjusted amplitude was different between stimulation locations (F[3,36]=11.935, p < .001), with the amplitude being significantly lower for sham relative to the other conditions, while the amplitude did not differ significantly between the active nVNS conditions cymba, Tragus and Cymba + Tragus (see Supplementary Tables 1 and 2). Participants then also rated the sensations of nVNS stimulation on the ear with VAS scales. The average intensity, pleasantness, burning, stinging, itching, tingling and vibration sensations are displayed in Figure 2. As intended, the perceived intensity did not differ between nVNS conditions, however, stronger stinging sensations were observed for the cymba (p=.031) and tragus (p=.011) relative to sham and also for the tragus relative to Cymba+tragus (p=.041).We note that the average stinging sensations remained below ~25% on the VAS scale. Other sensations associated with nVNS on the ear did not differ as a function of stimulation location (see Supplementary Table 4). Participants also rated their internal and other state sensations on VAS scales (see Figure 3) and none of these varied as a function of stimulation location at the start of the session (see Supplementary Table 7). These results show that nVNS stimulation location and contextual factors generally did not cause participants to feel any differently at the start of the experiment.

**Figure 2.**
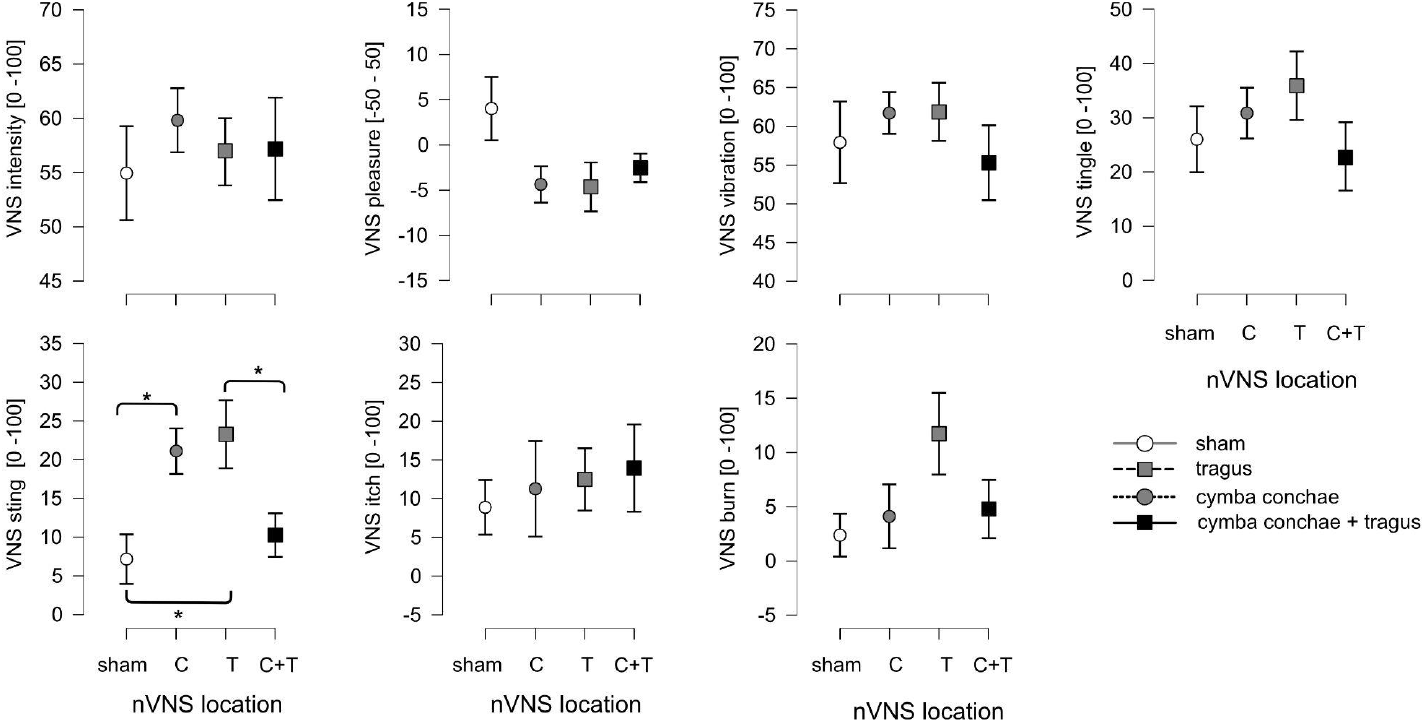
Participants sensations of nVNS stimulation on the ear. Graphs show average ratings ± standard error of the mean (SEM) across participants with respect to each nVNS stimulation location (on the x-axis). The nVNS stimulation locations are indicated with different symbols, open circles - sham/ear lobe, gray circles - cymba conchae, gray square - tragus, black square - cymba conchae and tragus. The different sensations that were rated on VAS scales are in separate graphs, labeled on the y-axis. Significant planned follow-up t-tests are indicated with an asterisk (P < 0.05, Bonferonni corrected for multiple comparisons).

**Figure 3.**
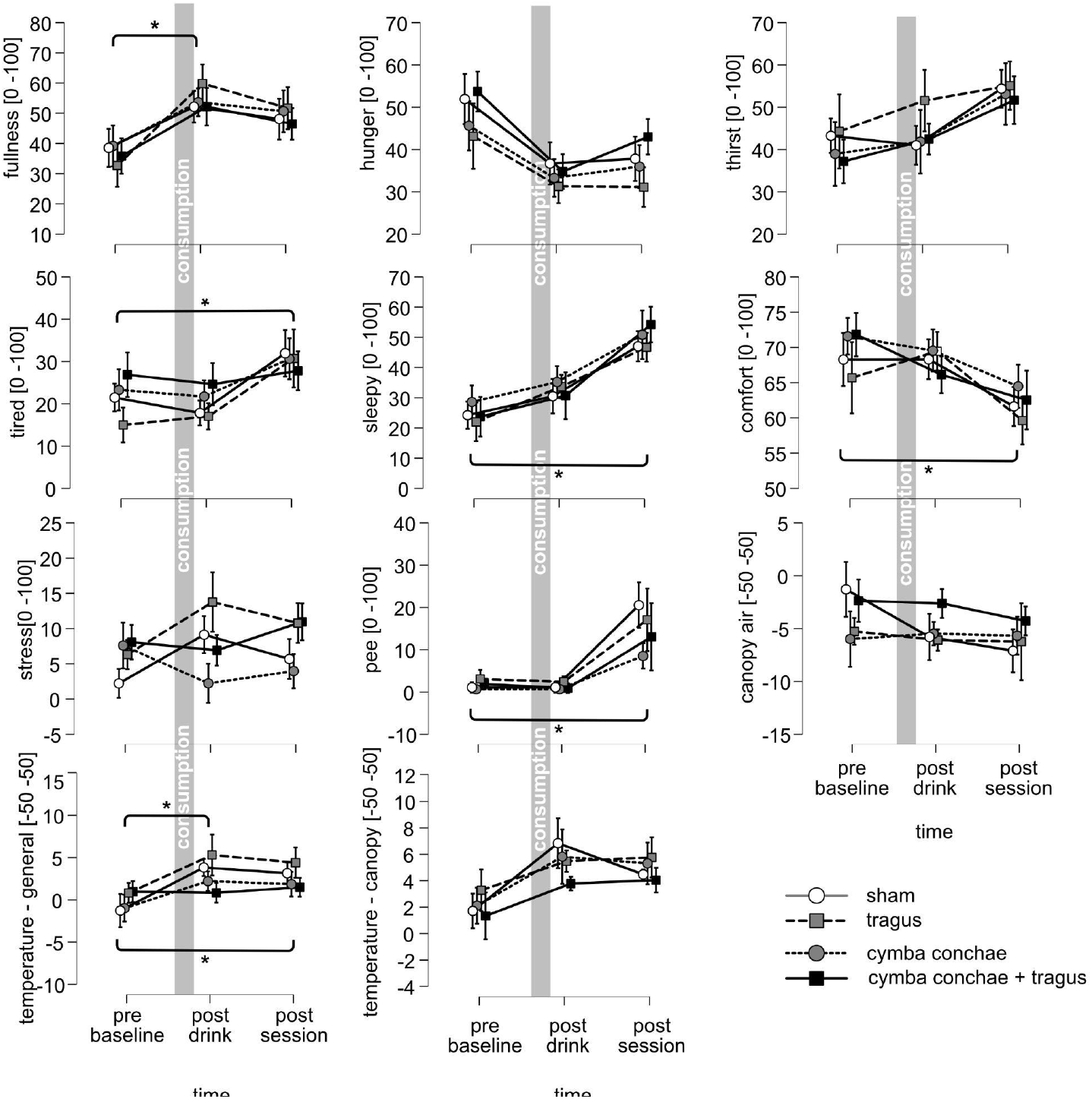
Participants sensations of internal and other states. Graphs show average ratings ± standard error of the mean (SEM) across participants per nVNS stimulation location. The nVNS stimulation locations are indicated with different symbols and lines, open circles + solid line - sham/ear lobe, gray circles + dotted line - cymba conchae, gray square + broken line - tragus, black square + solid line - cymba conchae and tragus. The x-axis reflects the factor time (pre-session, post-drink and post session). The different sensations that were rated on VAS scales are in separate graphs, labeled on the y-axis. Significant planned follow-up t-tests are indicated with an asterisk (P < 0.05, Bonferonni corrected for multiple comparisons).

### Acute nVNS in cymba conchae decreases heart-rate variability pre-drink consumption

Figure 4 shows the effects of acute nVNS in the different locations on the physiological measures of heart-rate variability and efferent metabolism over time, with the left panel of each graph showing the average (+/- standard error of the mean) across participants of the pre-drink consumption bin. To test whether acute nVNS affects the physiological measures, we examined the effect of nVNS location and bin (pre-drink, post-drink bins 1-7) on MGF, MGA rMSSD, and REE. We observed a significant effect of bin on MGA, rMSSD and REE, and a significant interaction of bin*location on rMSSD (Table 1). We observed no main effect of location on any of the physiological measures, nor a main effect of bin on MGF or an interaction of bin and location on MGF (Table 1). The planned t-test comparisons of location in the pre-drink bins for rMSSD (Table 2) show that there is a decreased heart-rate variability for acute nVNS in cymba conchae relative to lobe (p = .036) tragus (p = .033) and cymba conchae+tragus (p = .002). These results show that there are no differences in the effect of location of acute nVNS on REE and gastric frequency and gastric amplitude. However, nVNS in cymba conchae decreases heart-rate variability.

**Figure 4.**
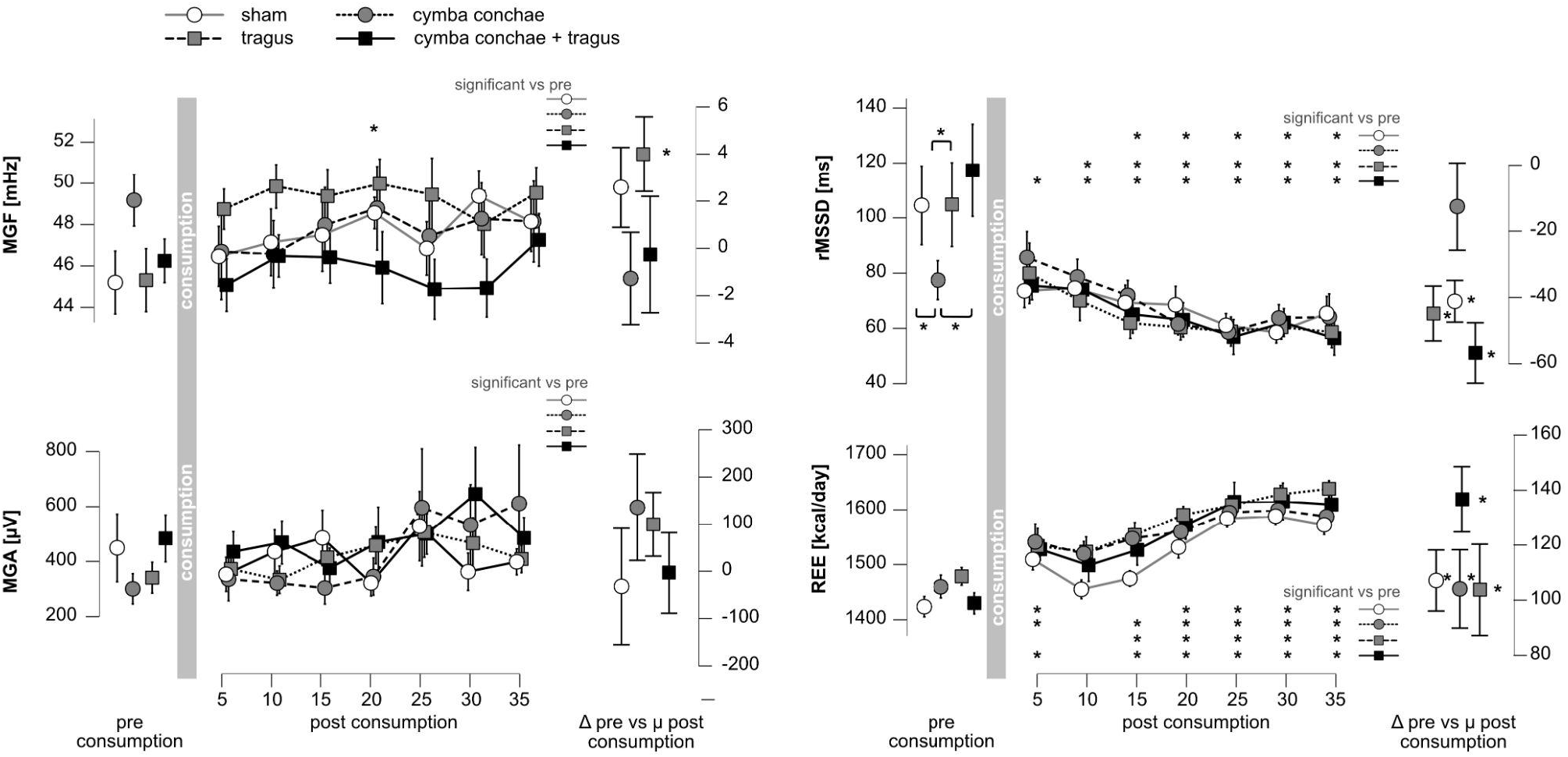
The effects of nVNS location on the physiological measurements. Graphs show average ratings ± standard error of the mean (SEM) across participants per nVNS stimulation locations, symbols and lines as in Figure 3. The x-axis reflects the factor bin (pre-drink, post-drink bins 1-7). The right panel reflects the change from pre-drink to the average of all post-drink bins. The top panel depicts raw mean gastric frequency (MGF in mHz), the center panel heart rate variability (rMSSD in ms) and the bottom panel resting energy expenditure (REE in kcal/day). Significant planned follow-up t-tests are indicated with an asterisk (P < 0.05, Bonferonni corrected for multiple comparisons). For the pre-consumption bin, significant differences between nVNS locations are indicated with an asterisk. In the post-consumption bins, the inset shows the time-bins that are significantly different relative to the pre-consumption time bin within nVNS location are indicated with an asterisk. For change from pre-drink to the average of all post-drink bins the asterisk indicates significant differences from 0 (one-sample t-test).

**Table 1.** Statistics of ANOVA of effects of bin and location of nVNS stimulation on physiology measures

| Measure | Effect of bin | Effect of location | Interaction bin* location |
| --- | --- | --- | --- |
| **MGF** | $F(7,91) = 1.837, p = .090, \eta^2 = 0.012$ | $F(3,39) = 0.833, p = .484, \eta^2 = 0.035$ | $F(21,273) = 1.205, p = .246, \eta^2 = 0.027$ |
| **MGA** | $F(7,91) = 2.334, p = .031, \eta^2 = 0.025$ | $F(3,39) = 0.299, p = .826, \eta^2 = 0.007$ | $F(21,273) = 0.811, p = .706, \eta^2 = 0.032$ |
| **rMSSD** | $F(7,91) = 5.887, p < .001, \eta^2 = 0.187$ | $F(3,39) = 0.100, p = 0.960, \eta^2 = 0.001$ | $F(21, 273) = 2.209, p = 0.002, \eta^2 = 0.038$ |
| **REE** | $F(7,91) = 33.35, p < .001, \eta^2 = 0.295$ | $F(3,39) = 1.343, p = .275, \eta^2 = 0.036$ | $F(21,273) = 0.880, p = .617, \eta^2 = 0.013$ |

**Table 2.**
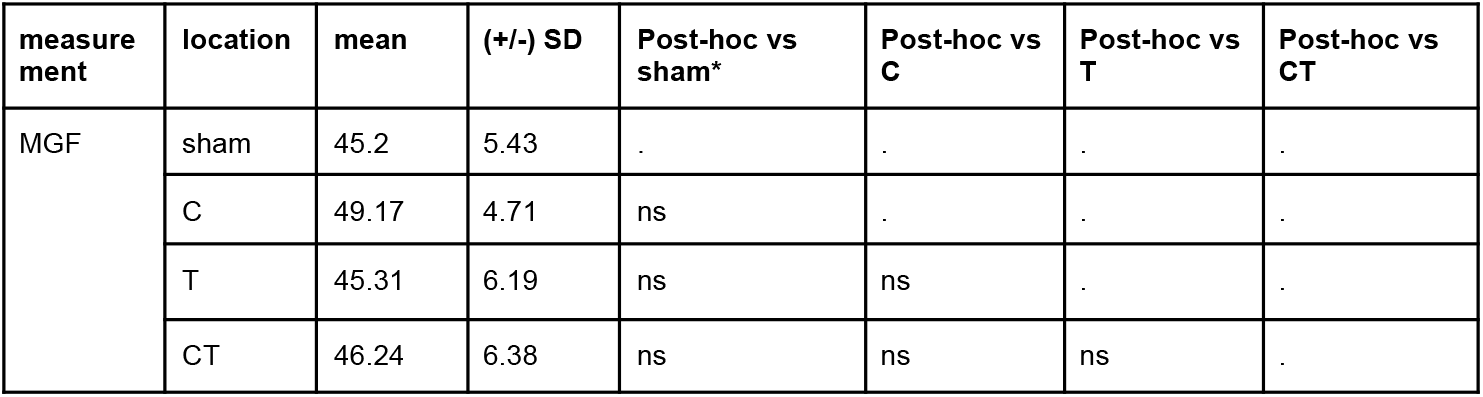
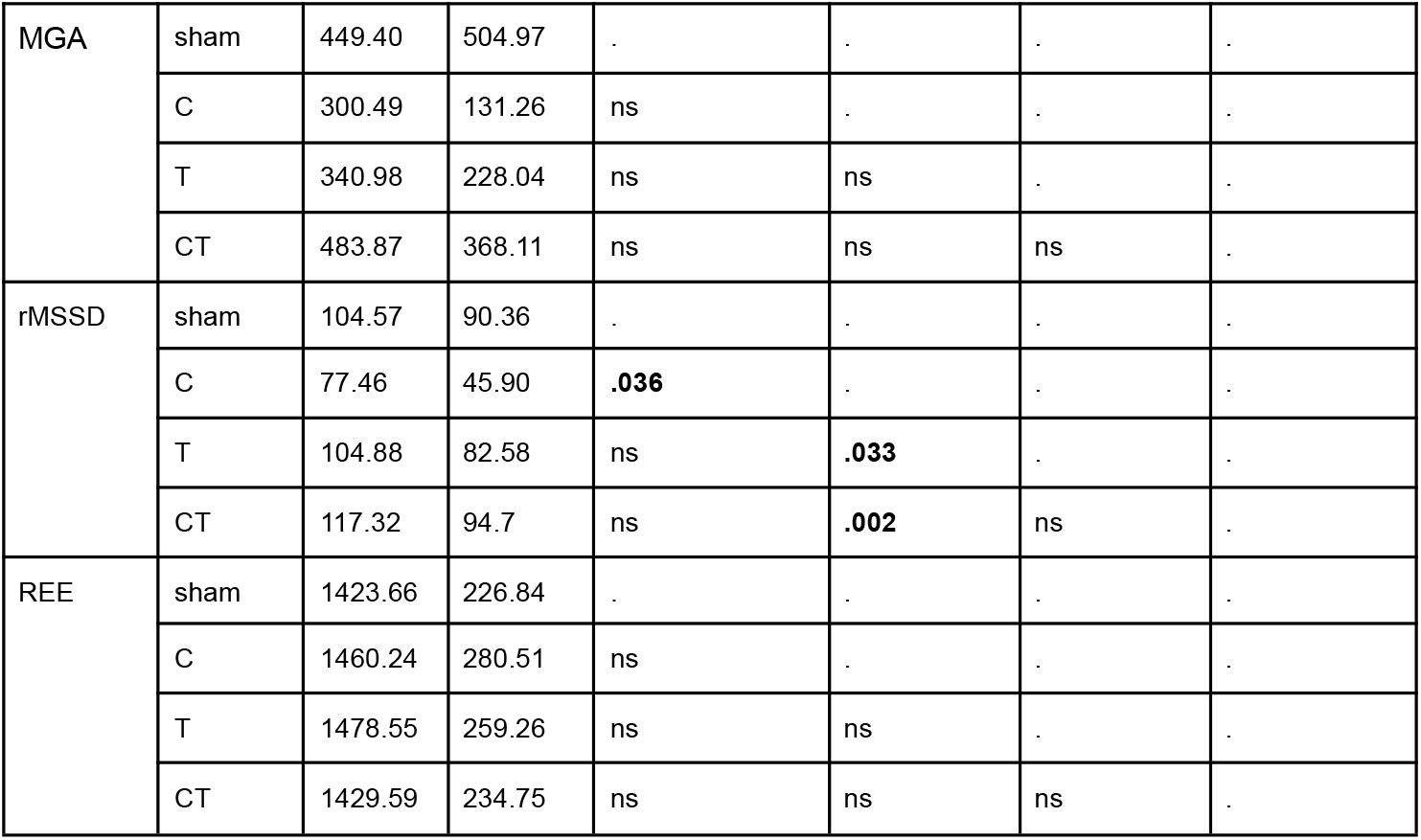
Descriptive statistics and planned comparison p-values of t-tests for effect of locations for the pre-drink baseline physiology measures

### The palatable drink is wanted less after nVNS in cymba conchae

After the baseline measurement of nVNS effects, participants were given a straw and instructed to take a sip of the palatable drink. They were then asked to rate their perception of the drink on VAS scales, see Figure 4 and Supplementary Table 10 for average (+/- standard error of the mean) across participants. Regardless of the nVNS location, participants generally like the drink and are motivated to consume it (ratings fall on average in the positive half of the bipolar VAS scales). To test for differences in perception and hedonics as a result of location of acute nVNS in the 15-minute pre-drink stimulation period, we examined the effect of nVNS location on each of the VAS scales. The drink is not perceived differently as a function of nVNS location, with the exception of a decreased motivation to consume the drink (“wanting”) after nVNS in cymba conchae relative to C+T (Supplementary Table 11).

After completing the ratings, participants consumed the remaining amount of the palatable drink. Then they rated their internal and other state sensations (see Figure 3). Participants were less hungry, and more full after they consumed the drink, regardless of the location of nVNS stimulation (see Supplementary Tables 5, 6). However, participants were not *differently* hungry, full or thirsty before they fully consume the palatable drink in different sessions with different nVNS stimulation locations (Supplementary Table 8), nor was there a difference in change from before consumption of the palatable drink to after consumption of the palatable drinks in different sessions (Supplementary Table 6).

These results show that acute nVNS in cymba conchae affects motivation to consume a drink, while other perceptual aspects of the drink, as well as internal state and other ratings are unaffected.

**Figure 5.**
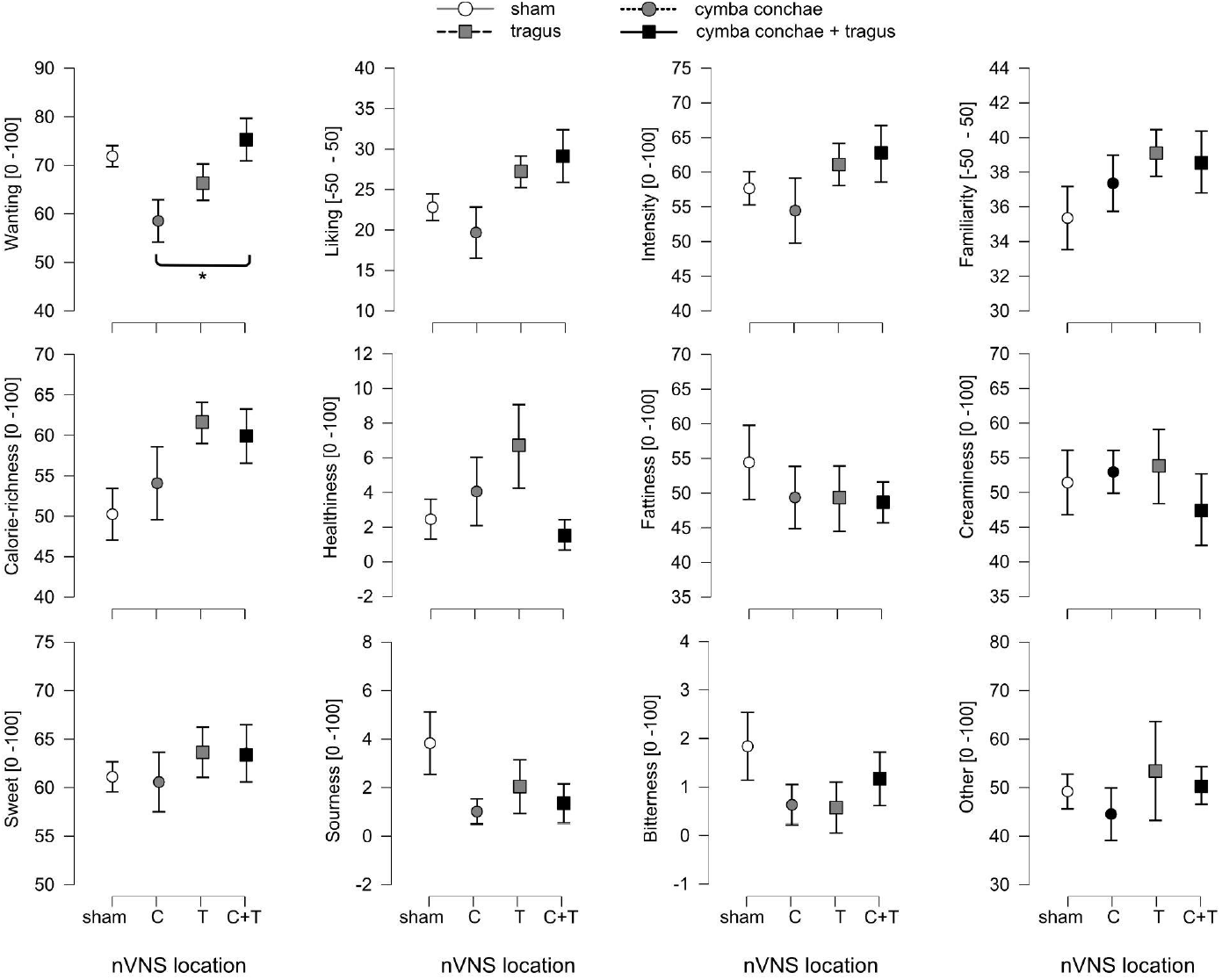
Participants’ perception of the palatable drink as a function of nVNS stimulation on the ear. Graphs show average ratings ± standard error of the mean (SEM) across participants per nVNS stimulation location (on the x-axis). The nVNS stimulation locations are indicated with different symbols, open circles - sham/ear lobe, gray circles - cymba conchae, gray square - tragus, black square - cymba conchae and tragus. The different sensations that were rated on VAS scales are in separate graphs, labeled on the y-axis. Significant planned follow-up t-tests are indicated with an asterisk (P < 0.05, Bonferonni corrected for multiple comparisons).

### After consumption of a palatable drink and with continued nVNS stimulation resting energy expenditure increases in all nVNS locations and heart-rate variability decreases in all nVNS locations except cymba conchae

The middle panel of each graph in Figure 2 shows the average (+/- standard error of the mean) of each physiological measure across participants in each of the post-drink consumption bins (up to 35 minutes). The right panel of each graph shows the change from pre-drink to the average of all post-drink bins. To test whether acute nVNS affects the physiological measures, we examined the effect of nVNS location and bin (pre-drink, post-drink bins 1-7) on rMSSD, MGF and DIT. We conducted planned follow-up t-test comparisons of the pre-drink bins relative to each of the post drink bins for each nVNS location. We also conducted planned follow-up one sample t-tests against 0 to test whether there was a change for the average of the post-consumption time bins relative to the pre-consumption bin. Regardless of location, we observed for both rMSSD and REE a change from pre- to post-drink bins (Table 1, right panel of Figure 2). For rMSSD, there was generally a decrease in post-drink consumption bins (i.e. decreased heart-rate variability) and for REE there was generally a robust increase in post-drink consumption bins (Table 3). For MGF, we observed no changes from pre- to post-drink consumption (no main effect of bin, Table 1), except for a small increase for nVNS in tragus (Table 3). For MGA we observed a main effect of bin. We note that this effect seems driven by a modest increase in the timepoints 20-35 minutes post-consumption (which was most pronounced for cymba conchae), but planned follow-up tests did not survive correction for multiple comparisons. We observed no main effect of nVNS location on any of the physiological measures (Table 1). However, we did observe an interaction between nVNS location and bin for rMSSD, such that rMSSD decreased for several bins of post-drink consumption in all nVNS locations *except* for cymba conchae (Table 3). In cymba conchae, rMSSD of the post-drink consumption bins were not significantly different from the pre-drink consumption bin.

**Table 3.** Planned comparison p-values for t-tests for separate post-consumption time bins relative to pre-consumption within location of nVNS stimulation

| measure<br>ment | locati<br>on | 1-5 min vs pre | | 6-10 min vs pre | | 11-15 min vs<br>pre | | 16-20 min vs<br>pre | | 21-25 min vs<br>pre | | 26-30 min vs pre | | 30-35 min vs pre | | $\Delta$ pre - $\mu$ post | |
| --- | --- | --- | --- | --- | --- | --- | --- | --- | --- | --- | --- | --- | --- | --- | --- | --- | --- |
|  |  | mean<br>(+/-sd) | P | mean<br>(+/-sd) | P | mean<br>(+/-sd) | P | mean<br>(+/-sd) | P | mean<br>(+/-sd) | P | mean<br>(+/-sd) | P | mean<br>(+/-sd) | P | T | p |
| MGF | sham | 46.46<br>(7.08) | ns | 47.14<br>(6.82) | ns | 47.49<br>(7.27) | ns | 48.55<br>(4.36) | ns | 46.84<br>(5.93) | ns | 49.38<br>(5.54) | ns | 48.14<br>(6.75) | ns | 1.385 | ns |
|  | C | 46.69<br>(9.79) | ns | 46.56<br>(6.99) | ns | 47.97<br>(7.43) | ns | 48.79<br>(8.08) | ns | 47.44<br>(9.37) | ns | 48.28<br>(7.60) | ns | 48.14<br>(8.42) | ns | -0.529 | ns |
|  | T | 48.74<br>(4.85) | ns | 49.85<br>(5.67) | ns | 49.37<br>(5.09) | ns | 49.97<br>(4.93) | **.047** | 49.44<br>(7.28) | ns | 48.02<br>(6.66) | ns | 49.53<br>(5.43) | ns | 2.583 | **0.023** |
|  | CT | 45.08<br>(5.63) | ns | 46.48<br>(4.86) | ns | 46.41<br>(5.70) | ns | 45.91<br>(6.30) | ns | 44.87<br>(6.00) | ns | 44.93<br>(5.85) | ns | 47.25<br>(4.83) | ns | -0.152 | ns |
| MGA | sham | 353.03<br>(180.61) | ns | 435.37<br>(335.14) | ns | 485.61<br>(431.82) | ns | 321.7<br>(210.48) | ns | 528.7<br>(496.89) | ns | 362.51<br>(219.2) | ns | 398.34<br>(259.77) | ns | -0.28 | ns |
|  | C | 335.49<br>(223.47) | ns | 321.22<br>(164.41) | ns | 303.51<br>(239.37) | ns | 344.94<br>(268.73) | ns | 596.5<br>(888.55) | ns | 532.87<br>(609.6) | ns | 611.52<br>(884.36) | ns | 1.258 | ns |
|  | T | 372.39<br>(189.51) | ns | 333.35<br>(194.1) | ns | 414.79<br>(310.21) | ns | 458.15<br>(334.28) | ns | 506.2<br>(376.27) | ns | 466.23<br>(380.55) | ns | 408.74<br>(251.4) | ns | 1.507 | ns |
|  | CT | 435.23<br>(329.62) | ns | 469.08<br>(396.99) | ns | 375.35<br>(340.4) | ns | 470.1<br>(500.74) | ns | 500.58<br>(328.52) | ns | 645.84<br>(667.78) | ns | 485.13<br>(315.4) | ns | -0.042 | ns |
| rMSSD | sham | 73.57<br>(64.30) | ns | 74.55<br>(58.24) | ns | 69.33<br>(57.12) | **.029** | 68.54<br>(49.96) | **.025** | 61.13<br>(42.94) | **.006** | 58.52<br>(39.88) | **.004** | 65.45<br>(41.53) | **.014** | -2.567 | **0.023** |
|  | C | 85.67<br>(73.27) | ns | 78.61<br>(63.41) | ns | 71.95<br>(55.99) | ns | 61.59<br>(42.40) | ns | 58.65<br>(42.39) | ns | 63.75<br>(35.16) | ns | 64.10<br>(39.35) | ns | -1.094 | ns |
|  | T | 79.93<br>(69.15) | ns | 70.03<br>(64.99) | **.032** | 61.91<br>(52.55) | **.007** | 60.37<br>(37.97) | **.005** | 58.86<br>(39.25) | **.004** | 60.45<br>(40.19) | **.005** | 58.64<br>(37.72) | **.004** | -2.804 | **0.015** |
|  | CT | 75.43<br>(56.06) | **.008** | 74.05<br>(56.70) | **.007** | 65.08<br>(46.83) | **.001** | 63.08<br>(42.69) | **.001** | 56.89<br>(37.65) | **<.001** | 62.10<br>(36.46) | **<.001** | 56.44<br>(33.38) | **<.001** | -2.724 | **0.017** |
| REE | sham | 1509.36<br>(238.41) | **.016** | 1455.06<br>(244.23) | ns | 1474.61<br>(224.82) | ns | 1531.78<br>(250.95) | **.003** | 1583.98<br>(221.51) | **<.001** | 1588.36<br>(237.07) | **<.001** | 1572.48<br>(236.97) | **<.001** | 7.529 | **<.001** |
|  | C | 1541.16<br>(300.27) | **.024** | 1519.95<br>(265.98) | ns | 1547.51<br>(269.69) | **.014** | 1559.87<br>(254.30) | **.005** | 1594.76<br>(288.86) | **<.001** | 1599.56<br>(260.75) | **<.001** | 1587.52<br>(257.12) | **<.001** | 8.261 | **<.001** |
|  | T | 1533.81<br>(336.37) | ns | 1523.34<br>(284.69) | ns | 1554.02<br>(287.71) | **.039** | 1590.61<br>(277.41) | **.002** | 1608.27<br>(271.79) | **<.001** | 1627.87<br>(262.06) | **<.001** | 1638.11<br>(266.16) | **<.001** | 5.884 | **<.001** |
|  | CT | 1527.74<br>(292.63) | **.006** | 1498.59<br>(270.47) | ns | 1526.58<br>(274.89) | **.006** | 1571.97<br>(269.60) | **.003** | 1614.35<br>(266.68) | **<.001** | 1615.29<br>(257.57) | **<.001** | 1609.44<br>(250.92) | **<.001** | 6.718 | **<.001** |

To assess the role of changes in internal and other states, we examined planned follow-up t-tests on VAS ratings and observed that participants were not *differently* hungry, full or thirsty (or anything else) at the end of the test-session with respect to different nVNS stimulation locations (Supplementary Table 9).

These results show that nVNS in cymba conchae does not lead to a decrease in heart-rate variability after consumption of a drink, while the other locations including the control condition do show a decrease after consumption.

## Discussions

The aim of this study was to compare the effect of different stimulation locations in the outer ear for nVNS during hungry and full states on proxies for autonomic outflow and efferent metabolism: heart-rate variability, gastric frequency and amplitude, and REE. In agreement with our first prediction, we observed that nVNS in cymba conchae (but not in tragus and the combination of cymba conchae + tragus) modulated autonomic outflow as expressed by heart-rate variability. More specifically, nVNS in cymba conchae decreased heart-rate variability in a hungry state. Additionally, while heart-rate variability decreased in the other stimulation locations post-drink consumption, it did not change from pre- to post-drink consumption with nVNS in the cymba conchae location. REE and gastric frequency did not show effects of stimulation location. Gastric amplitude showed a trend to increase late postprandially with nVNS in the cymba conchae.

Fasting increases heart-rate variability, and food consumption decreases heart-rate variability [41]. As we observed a pre-drink decrease with nVNS in cymba conchae and a lack of a post-drink decrease in heart-rate variability (which was consistently observed in the sham and other conditions), we suggest that nVNS in cymba conchae affects heart-rate variability in a similar manner as food consumption does. Intriguingly, we also observed a decrease in pre-drink motivation to consume the drink (“wanting”) after nVNS in cymba conchae. This is in agreement with the observation that nVNS increases drive to obtain less-wanted prospective food rewards [22] and increases liking of less-liked low-fat puddings [21]. Other studies have not shown effects of increases in liking or wanting of food (images) in healthy controls [42,43]. Perhaps effects of nVNS vary as a function of food stimulus saliency, which is presumably higher when it is sufficiently proximal (in the mouth) [21] or the object of a goal-directed task

[22]. Taken together, we speculate that nVNS may modulate primarily motivation for a proximal food reward. Future studies may focus on comparing effects of nVNS on orally sampled foods relative to food images.

We observed a robust increase in resting energy expenditure post drink-consumption, which is the expected thermic effect of food. However, no interaction with the location of nVNS was observed. In previous reports, a lack of effect of nVNS on REE (in the absence of food consumption [24], and a lack of an effect 160 minutes post glucose-drink consumption [25] have been observed. Our current results extend these previous findings to the time-frame of the thermic effects of food in the cephalic and early phases of digestion (0-35 minutes post consumption).

We observed no effect of drink consumption or location of nVNS on gastric frequency. Gastric frequency is generally relatively unaffected by food consumption [44], however a previous report showed that nVNS in cymba conchae reduces gastric frequency in a fasted resting state [24]. We do not replicate this effect, although our sample size may have been too small to observe subtle effects. Gastric amplitude generally increases postprandially in healthy participants [44]. We observed a small increase in gastric amplitude post-drink consumption (in later portions of the measurement) with nVNS in cymba conchae.The direction of this effect is in agreement with the effect on heart-rate variability and wanting; an effect that is normally observed with food-consumption is induced or enhanced with nVNS in cymba conchae. As gastric activity is depressed with sleep [45], it is possible that the increased sleepiness of participants towards the end of the sessions decreased our ability to observe effects on electrogastrogram. We recommend the use of entertainment for the participant in the form of low-arousal media consumption, such as listening to an audio-book or watching a documentary, to reduce sleepiness and increase sensitivity to gastric activity in future postprandial studies.

Our second prediction was that if the vagally innervated areas of the outer ear in cymba conchae and tragus are functionally equivalent, we should observe that stimulation of the combination of cymba conchae + tragus leads to effects in the same *direction* (relative to sham in ear lobe) and of a *magnitude* that is equal or greater than the separate locations in cymba conchae or tragus, but not smaller. We did not consistently observe differences for the combined location (as well as tragus alone) relative to earlobe. When taken together with mixed directions of the effect relative to earlobe, we find inconclusive evidence towards the equivalence of the vagally innervated areas of tragus and cymba conchae. This also indicates there may be no advantage to stimulating both locations simultaneously, at least not when the goal is to affect outcomes related to food reward and metabolism.

There are a few limitations to this study. We had a relatively small sample size, which was sufficient to detect robust effects of for example postprandial increases in REE, but it is possible some more subtle effects were missed. It is unclear if the stimulation parameters used here (chosen by convention of use in most studies in the literature) are optimal for stimulation of the vagus nerve and for affecting outcomes related to food reward and metabolism. The palatable drink was always consumed after about 20 minutes of baseline nVNS stimulation, so it is not clear how nVNS onset simultaneously with onset of drink consumption would affect the variables measured here. While we optimized the study design for a long post-drink measurement, we could not measure the full intended 45-60 minutes for all participants. We asked participants to void their bladder immediately before the session and we offered a comfortable bed and continued monitoring comfort throughout the session. However, some sessions were not completed for the full duration due to an urgent need to urinate. Also, while participants were comfortable, they also grew sleepy, which may have negatively affected our ability to measure electrogastrogram.

In sum, we show that nVNS -- in cymba conchae only -- affects autonomic outflow as reflected by heart-rate variability in a hungry state, which was accompanied by a decrease in wanting of a prospective palatable drink. After consumption of the drink, nVNS in the other stimulation conditions (including sham stimulation) show the canonical post-prandial decrease in heart-rate variability, which is absent for nVNS in the cymba conchae. With nVNS in cymba conchae we also observed a trend for increased gastric amplitude in the late time-points post-drink consumption. We observed no effects of nVNS location on gastric frequency or resting energy expenditure. The direction of effects of tVNS in cymba conchae are in agreement, in that they are all normally observed with food-consumption, and that in the current paper they each were induced (pre-consumption) or enhanced (post-consumption) with nVNS in cymba conchae. We conclude that nVNS in cymba conchae may act primarily on vagal afferent autonomic and only modestly on metabolic outputs in a similar way as food consumption does. nVNS may thus specifically continue to be explored for treating overeating and/or supporting diet-based weight-loss programs.

## Supplementary Materials

**Supplementary Table 1.** Descriptive statistics of nVNS stimulation in amplitude expressed as 0-100% of 20 mA per location

|  | **Sham** | **C** | **T** | **CT** |
| --- | --- | --- | --- | --- |
|  | **mean (+/-sd)** | **mean (+/-sd)** | **mean (+/-sd)** | **mean (+/-sd)** |
| Percentage of 20 mA | 13.96 (4.19) | 35.85 (18.85) | 34.15 (14.24) | 25.89 (10.47) |

**Supplementary Table 1.** Planned comparison t-tests for effect of location

|  | **Sham** | **C** | **T** | **CT** |
| --- | --- | --- | --- | --- |
| **Sham** | . | . | . | . |
| **C** | < .001 | . | . | . |
| **T** | < .001 | ns | ns |  |
| **CT** | 0.024 | ns | ns | ns |

\*Bonferroni corrected for 6 comparisons

**Supplementary Table 3.** Descriptive statistics of perceptual ratings of nVNS stimulation per location

| **VAS label** | **Sham** | **C** | **T** | **CT** |
| --- | --- | --- | --- | --- |
|  | **mean (+/-sd)** | **mean (+/-sd)** | **mean (+/-sd)** | **mean (+/-sd)** |
| Intensity | 54.95 (21.21) | 59.82 (19.49) | 56.91 (9.42) | 57.18 (19.19) |
| Pleasantness | 4.04 (8.11) | -4.36 (11.58) | -4.64 (14.02) | -2.55 (8.01) |
| Sting | 7.17 (18.25) | 21.11 (18.75) | 23.29 (25.38) | 10.26 (21.52) |
| Burn | 2.38 (5.39) | 4.12 (4.78) | 11.73 (20.96) | 4.79 (15.94) |
| Tingle | 26.03 (29.47) | 30.85 (25.87) | 35.9 (26.56) | 22.87 (17.76) |
| Itching | 8.89 (18.79) | 11.28 (16.65) | 12.49 (20.31) | 13.96 (26.67) |
| Vibration | 57.92 (25.21) | 61.71 (22.32) | 61.86 (17.24) | 55.29 (25.37) |

**Supplementary Table 4.**
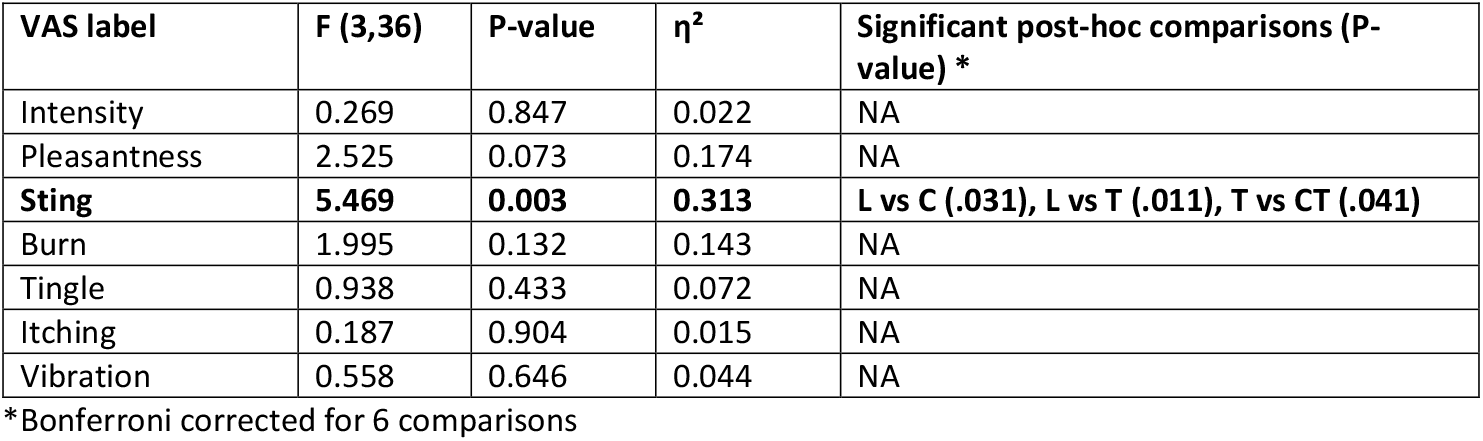
Statistics of ANOVA of effect of location on perceptual ratings of nVNS stimulation

| **VAS label** | **F (3,36)** | **P-value** | **$\eta^2$** | **Significant post-hoc comparisons (P-value) \*** |
| --- | --- | --- | --- | --- |
| Intensity | 0.269 | 0.847 | 0.022 | NA |
| Pleasantness | 2.525 | 0.073 | 0.174 | NA |
| **Sting** | **5.469** | **0.003** | **0.313** | **L vs C (.031), L vs T (.011), T vs CT (.041)** |
| Burn | 1.995 | 0.132 | 0.143 | NA |
| Tingle | 0.938 | 0.433 | 0.072 | NA |
| Itching | 0.187 | 0.904 | 0.015 | NA |
| Vibration | 0.558 | 0.646 | 0.044 | NA |

\*Bonferroni corrected for 6 comparisons

**Supplementary Table 5.** Statistics of ANOVA of internal and other state ratings

| Scale | Effect of time | Effect of location | Interaction time*location |
| --- | --- | --- | --- |
| Hunger | **F(2,22) = 3.006, p = 0.07, $\eta^2$ = 0.123** | F(3,33) = 1.713, p = 0.183, $\eta^2$ = 0.035 | F(6,66) = 0.5, p = 0.806, $\eta^2$ = 0.007 |
| Fullness | **F(2,22) = 4.566, p = 0.022, $\eta^2$ = 0.124** | F(3,33) = 0.121, p = 0.947, $\eta^2$ = 0.004 | F(6,66) = 0.571, p = 0.752, $\eta^2$ = 0.011 |
| Thirst | F(2,22) = 2.316, p = 0.122, $\eta^2$ = 0.068 | F(3,33) = 0.476, p = 0.701, $\eta^2$ = 0.015 | F(6,66) = 0.344, p = 0.911, $\eta^2$ = 0.008 |
| Need to pee | **F(2,22) = 21.325, p = < .001, $\eta^2$ = 0.208** | F(3,33) = 0.894, p = 0.455, $\eta^2$ = 0.017 | F(6,66) = 0.524, p = 0.788, $\eta^2$ = 0.021 |
| Tiredness | **F(2,22) = 4.938, p = 0.017, $\eta^2$ = 0.079** | F(3,33) = 0.408, p = 0.748, $\eta^2$ = 0.017 | F(6,66) = 1.085, p = 0.381, $\eta^2$ = 0.024 |
| Sleepiness | **F(2,22) = 9.299, p = 0.001, $\eta^2$ = 0.234** | F(3,33) = 0.313, p = 0.816, $\eta^2$ = 0.007 | F(6,66) = 0.328, p = 0.92, $\eta^2$ = 0.007 |
| Stress | F(2,22) = 1.694, p = 0.207, $\eta^2$ = 0.009 | F(3,33) = 1.412, p = 0.257, $\eta^2$ = 0.059 | F(6,66) = 1.937, p = 0.088, $\eta^2$ = 0.063 |
| Comfort | **F(2,22) = 4.457, p = 0.024, $\eta^2$ = 0.083** | F(3,33) = 0.525, p = 0.668, $\eta^2$ = 0.014 | F(6,66) = 0.454, p = 0.84, $\eta^2$ = 0.016 |
| Temperature | **F(2,22) = 4.842, p = 0.018, $\eta^2$ = 0.076** | F(3,33) = 1.472, p = 0.24, $\eta^2$ = 0.038 | F(6,66) = 0.745, p = 0.615, $\eta^2$ = 0.027 |
| Canopy temperature | F(2,22) = 2.667, p = 0.092, $\eta^2$ = 0.095 | F(3,33) = 1.109, p = 0.359, $\eta^2$ = 0.019 | F(6,66) = 0.473, p = 0.826, $\eta^2$ = 0.012 |
| Canopy air | F(2,22) = 0.376, p = 0.691, $\eta^2$ = 0.016 | F(3,33) = 1.132, p = 0.35, $\eta^2$ = 0.027 | F(6,66) = 1.291, p = 0.273, $\eta^2$ = 0.023 |

**Supplementary Table 6.** Planned comparison p-values for t-tests for interaction time and location of nVNS on internal and other state ratings

| VAS label | Location | Pre-session vs post drink* | pre-session vs post session* | $\Delta$ pre vs post drink** | | | |
| --- | --- | --- | --- | --- | --- | --- | --- |
|  |  |  |  | Post-hoc vs sham* | Post-hoc vs C | Post-hoc vs T | Post-hoc vs CT |
| Hunger | All locations | ns | ns |  |  |  |  |
|  | Sham | ns | ns | . | . | . | . |
|  | C | ns | ns | ns | . | . | . |
|  | T | ns | ns | ns | ns | . | . |
|  | CT | 0.045 | ns | ns | ns | ns | . |
| **Fullness** | All locations | .023 | ns |  |  |  |  |
|  | Sham | ns | ns | . | . | . | . |
|  | C | ns | ns | ns | . | . | . |
|  | T | 0.010 | ns | ns | ns | . | . |
|  | CT | ns | ns | ns | ns | ns | . |
| **Thirst** | All locations | ns | ns |  |  |  |  |
|  | Sham | ns | ns | . | . | . | . |
|  | C | ns | ns | ns | . | . | . |
|  | T | ns | ns | ns | ns | . | . |
|  | CT | ns | ns | ns | ns | ns | . |
| **Need to pee** | All locations | ns | < .001 |  |  |  |  |
|  | Sham | ns | 0.006 | . | . | . | . |
|  | C | ns | ns | ns | . | . | . |
|  | T | ns | 0.042 | ns | ns | . | . |
|  | CT | ns | ns | ns | ns | ns | . |
| **Tiredness** | All locations | ns | .04 |  |  |  |  |
|  | Sham | ns | ns | . | . | . | . |
|  | C | ns | ns | ns | . | . | . |
|  | T | ns | 0.022 | ns | ns | . | . |
|  | CT | ns | ns | ns | ns | ns | . |
| **Sleepiness** | All locations | ns | 0.001 |  |  |  |  |
|  | Sham | ns | 0.030 | . | . | . | . |
|  | C | ns | ns | ns | . | . | . |
|  | T | ns | 0.019 | ns | ns | . | . |
|  | CT | ns | 0.015 | ns | ns | ns | . |
| **Stress** | All locations | ns | ns |  |  |  |  |
|  | Sham | ns | ns | . | . | . | . |
|  | C | ns | ns | ns | . | . | . |
|  | T | ns | ns | ns | ns | . | . |
|  | CT | ns | ns | ns | ns | ns | . |
| **Comfort** | All locations | ns | 0.035 |  |  |  |  |
|  | Sham | ns | ns | . | . | . | . |
|  | C | ns | ns | ns | . | . | . |
|  | T | ns | ns | ns | ns | . | . |
|  | CT | ns | ns | ns | ns | ns | . |
| **Temperature** | All locations | 0.029 | 0.037 |  |  |  |  |
|  | Sham | ns | ns | . | . | . | . |
|  | C | ns | ns | ns | . | . | . |
|  | T | ns | ns | ns | ns | . | . |
|  | CT | ns | ns | ns | ns | ns | . |
| **Canopy temperature** | All locations | ns | ns |  |  |  |  |
|  | Sham | ns | ns | . | . | . | . |
|  | C | ns | ns | ns | . | . | . |
|  | T | ns | ns | ns | ns | . | . |
|  | CT | ns | ns | ns | ns | ns | . |
| **Canopy air** | All locations | ns | ns |  |  |  |  |
|  | Sham | ns | ns | . | . | . | . |
|  | C | ns | ns | ns | . | . | . |
|  | T | ns | ns | ns | ns | . | . |
|  | CT | ns | ns | ns | ns | ns | . |

\*Bonferroni corrected for 2 comparisons
\*\*Bonferroni corrected for 6 comparisons

**Supplementary Table 7.** Planned comparison p-values for t-tests for effect of location of nVNS on internal and other state ratings at the pre-baseline time-point only

| VAS label | Location | Pre baseline* |  |  |  |
| --- | --- | --- | --- | --- | --- |
|  |  | T-test vs sham* | T-test vs C | T-test vs T | T-test vs CT |
| **Hunger** | Sham | . | . | . | . |
|  | C | ns | . | . | . |
|  | T | ns | ns | . | . |
|  | CT | ns | ns | ns | . |
| **Fullness** | Sham | . | . | . | . |
|  | C | ns | . | . | . |
|  | T | ns | ns | . | . |
|  | CT | ns | ns | ns | . |
| **Thirst** | Sham | . | . | . | . |
|  | C | ns | . | . | . |
|  | T | ns | ns | . | . |
|  | CT | ns | ns | ns | . |
| **Need to pee** | Sham | . | . | . | . |
|  | C | ns | . | . | . |
|  | T | ns | ns | . | . |
|  | CT | ns | ns | ns | . |
| **Tiredness** | Sham | . | . | . | . |
|  | C | ns | . | . | . |
|  | T | ns | ns | . | . |
|  | CT | ns | ns | ns | . |
| **Sleepiness** | Sham | . | . | . | . |
|  | C | ns | . | . | . |
|  | T | ns | ns | . | . |
|  | CT | ns | ns | ns | . |
| **Stress** | Sham | . | . | . | . |
|  | C | ns | . | . | . |
|  | T | ns | ns | . | . |
|  | CT | ns | ns | ns | . |
| **Comfort** | Sham | . | . | . | . |
|  | C | ns | . | . | . |
|  | T | ns | ns | . | . |
|  | CT | ns | ns | ns | . |
| **Temperature** | Sham | . | . | . | . |
|  | C | ns | . | . | . |
|  | T | ns | ns | . | . |
|  | CT | ns | ns | ns | . |
| **Canopy temperature** | Sham | . | . | . | . |
|  | C | ns | . | . | . |
|  | T | ns | ns | . | . |
|  | CT | ns | ns | ns | . |
| **Canopy air** | Sham | . | . | . | . |
|  | C | ns | . | . | . |
|  | T | ns | ns | . | . |
|  | CT | ns | ns | ns | . |

\*Bonferroni corrected for 6 comparisons

**Supplementary Table 8.** Planned comparison p-values for t-tests for effect of location of nVNS on internal and other state ratings at the post drink-consumption time-point only

| VAS label | Location | Post drink-consumption* |  |  |  |
| --- | --- | --- | --- | --- | --- |
|  |  | T-test vs sham | T-test vs C | T-test vs T | T-test vs CT |
| **Hunger** | Sham | . | . | . | . |
|  | C | ns | . | . | . |
|  | T | ns | ns | . | . |
|  | CT | ns | ns | ns | . |
| **Fullness** | Sham | . | . | . | . |
|  | C | ns | . | . | . |
|  | T | ns | ns | . | . |
|  | CT | ns | ns | ns | . |
| **Thirst** | Sham | . | . | . | . |
|  | C | ns | . | . | . |
|  | T | ns | ns | . | . |
|  | CT | ns | ns | ns | . |
| **Need to pee** | Sham | . | . | . | . |
|  | C | ns | . | . | . |
|  | T | ns | ns | . | . |
|  | CT | ns | ns | ns | . |
| **Tiredness** | Sham | . | . | . | . |
|  | C | ns | . | . | . |
|  | T | ns | ns | . | . |
|  | CT | ns | ns | ns | . |
| **Sleepiness** | Sham | . | . | . | . |
|  | C | ns | . | . | . |
|  | T | ns | ns | . | . |
|  | CT | ns | ns | ns | . |
| **Stress** | Sham | . | . | . | . |
|  | C | ns | . | . | . |
|  | T | ns | ns | . | . |
|  | CT | ns | ns | ns | . |
| **Comfort** | Sham | . | . | . | . |
|  | C | ns | . | . | . |
|  | T | ns | ns | . | . |
|  | CT | ns | ns | ns | . |
| **Temperature** | Sham | . | . | . | . |
|  | C | ns | . | . | . |
|  | T | ns | ns | . | . |
|  | CT | ns | ns | ns | . |
| **Canopy temperature** | Sham | . | . | . | . |
|  | C | ns | . | . | . |
|  | T | ns | ns | . | . |
|  | CT | ns | ns | ns | . |
| **Canopy air** | Sham | . | . | . | . |
|  | C | ns | . | . | . |
|  | T | ns | ns | . | . |
|  | CT | ns | ns | ns | . |

\*Bonferroni corrected for 6 comparisons

**Supplementary Table 9.** Planned comparison p-values for t-tests for effect of location of nVNS on internal and other state ratings at the post session time-point only

| VAS label | Location | Post session* |  |  |  |
| --- | --- | --- | --- | --- | --- |
|  |  | T-test vs sham | T-test vs C | T-test vs T | T-test vs CT |
| **Hunger** | Sham | . | . | . | . |
|  | C | ns | . | . | . |
|  | T | ns | ns | . | . |
|  | CT | ns | ns | ns | . |
| **Fullness** | Sham | . | . | . | . |
|  | C | ns | . | . | . |
|  | T | ns | ns | . | . |
|  | CT | ns | ns | ns | . |
| **Thirst** | Sham | . | . | . | . |
|  | C | ns | . | . | . |
|  | T | ns | ns | . | . |
|  | CT | ns | ns | ns | . |
| **Need to pee** | Sham | . | . | . | . |
|  | C | ns | . | . | . |
|  | T | ns | ns | . | . |
|  | CT | ns | ns | ns | . |
| **Tiredness** | Sham | . | . | . | . |
|  | C | ns | . | . | . |
|  | T | ns | ns | . | . |
|  | CT | ns | ns | ns | . |
| **Sleepiness** | Sham | . | . | . | . |
|  | C | ns | . | . | . |
|  | T | ns | ns | . | . |
|  | CT | ns | ns | ns | . |
| **Stress** | Sham | . | . | . | . |
|  | C | ns | . | . | . |
|  | T | ns | ns | . | . |
|  | CT | ns | ns | ns | . |
| **Comfort** | Sham | . | . | . | . |
|  | C | ns | . | . | . |
|  | T | ns | ns | . | . |
|  | CT | ns | ns | ns | . |
| **Temperature** | Sham | . | . | . | . |
|  | C | ns | . | . | . |
|  | T | ns | ns | . | . |
|  | CT | ns | ns | ns | . |
| **Canopy temperature** | Sham | . | . | . | . |
|  | C | ns | . | . | . |
|  | T | ns | ns | . | . |
|  | CT | ns | ns | ns | . |
| **Canopy air** | Sham | . | . | . | . |
|  | C | ns | . | . | . |
|  | T | ns | ns | . | . |
|  | CT | ns | ns | ns | . |

\*Bonferroni corrected for 6 comparisons

**Supplementary Table 10.**
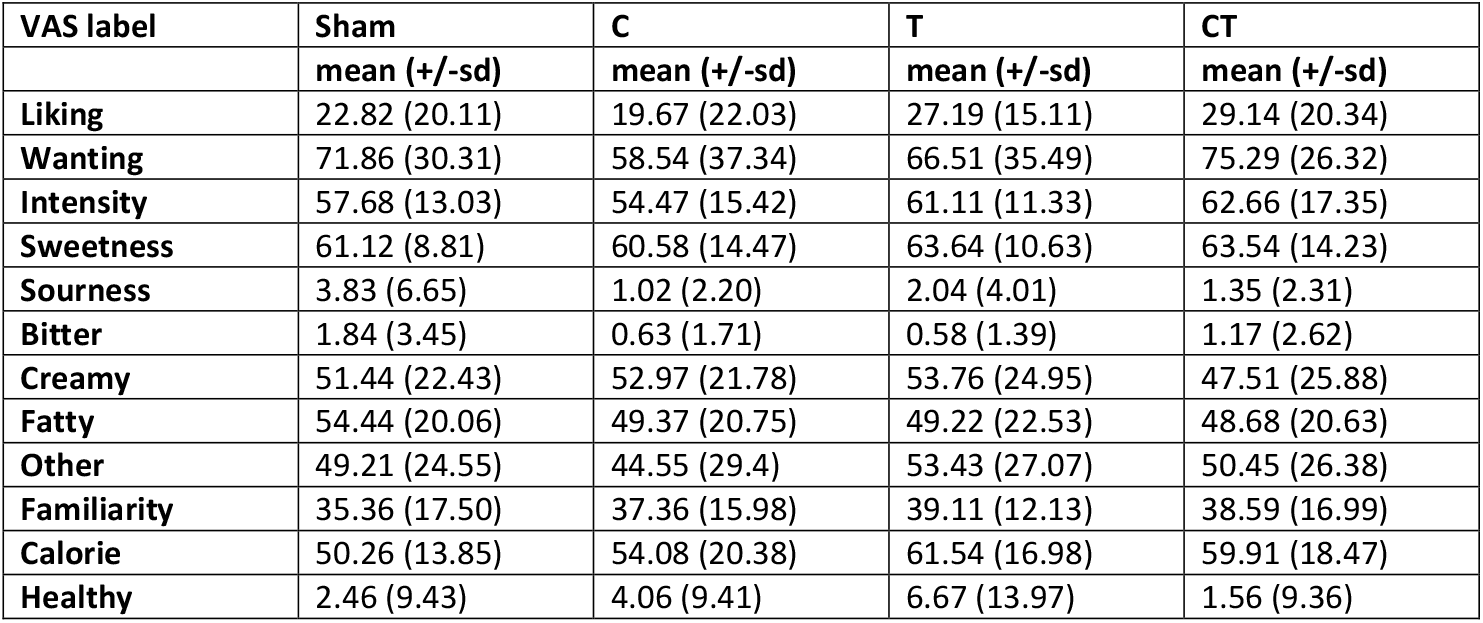
Descriptive statistics of perceptual ratings of the palatable drink per location

**Supplementary Table 11.** Statistics of ANOVA of perceptual ratings of the palatable drink per location

| VAS label | F(3,33) | p | $\eta^2$ | Significant post-hoc comparisons (P-value) * |
| --- | --- | --- | --- | --- |
| Liking | 2.69 | 0.062 | 0.196 | NA |
| Wanting | **3.72** | **0.021** | **0.253** | **C vs CT (.022)** |
| Intensity | 1.004 | 0.403 | 0.084 | NA |
| Sweetness | 0.375 | 0.772 | 0.033 | NA |
| Sourness | 1.669 | 0.193 | 0.132 | NA |
| Bitter | 1.109 | 0.359 | 0.092 | NA |
| Creamy | 0.358 | 0.784 | 0.031 | NA |
| Fatty | 0.365 | 0.779 | 0.032 | NA |
| Other | 0.339 | 0.797 | 0.03 | NA |
| Familiarity | 1.019 | 0.397 | 0.085 | NA |
| Calorie | 2.261 | 0.100 | 0.170 | NA |
| Healthy | 1.702 | 0.186 | 0.134 | NA |

\*Bonferroni corrected for 6 comparisons

